# The null additivity of multi-drug combinations

**DOI:** 10.1101/239517

**Authors:** D. Russ, R. Kishony

## Abstract

From natural ecology ^1–4^ to clinical therapy ^5–8^, cells are often exposed to mixtures of multiple drugs. Two competing null models are used to predict the combined effect of drugs: response additivity (Bliss) and dosage additivity (Loewe) ^9–11^. Here, noting that these models diverge with increased number of drugs, we contrast their predictions with measurements of *Escherichia coli* growth under combinations of up to 10 different antibiotics. As the number of drugs increases, Bliss maintains accuracy while Loewe systematically loses its predictive power. The total dosage required for growth inhibition, which Loewe predicts should be fixed, steadily increases with the number of drugs, following a square root scaling. This scaling is explained by an approximation to Bliss where, inspired by RA Fisher’s classical geometric model ^12^, dosages of independent drugs adds up as orthogonal vectors rather than linearly. This dose-orthogonality approximation provides results similar to Bliss, yet uses the dosage language as Loewe and is hence easier to implement and intuit. The rejection of dosage additivity in favor of effect additivity and dosage orthogonality provides a framework for understanding how multiple drugs and stressors add up in nature and the clinic.

## Main Text

In both nature and the clinic, cells are often exposed to combinations of multiple stresses and drugs. In natural ecosystems, such as the soil, dozens of microbial species capable of producing different antimicrobial compounds coexist in very close proximity, thus exposing each-others to a mixture of multiple stressors ^1–4^ In clinical settings, drug combinations, aimed at reducing side effects and counteracting resistance ^13–17^, are becoming increasingly important in treatment for infectious diseases and cancer ^5–8, 18, 19^. It is therefore of wide importance to understand how cell growth is affected by combinations of multitude of stressors and what are thereby the general rules of high-dimensionality drug arithmetics.

When combined together, drugs can interact to synergize or antagonize each other effects relative to a null additive model. Synergy occurs when the combined effect of drugs is larger than expected based on their individual effects. Conversely, drugs can also antagonize each other, leading to a combined effect that is smaller than expected. These interactions are important in clinical settings as a way of increasing treatment potency and selectivity ^20–22^ or slowing selection for resistance ^14,17,23^. Importantly, the definition of both synergy and antagonism relies on comparing the effect of drug combinations with a null model of “additive expectation” ^6,24–27^.

There are two primary models for the null effect of drug combinations ^6,28^: the Bliss model ^10,29^, which assumes response additivity and the Loewe model ^9,11^, which assumes dose additivity. According to Bliss, the combined effect of two drugs *E*_1+2_ is simply the sum of their individual effects ^30^, *E*_1+2_ = *E_1_ + E*_2_, where *E, = (g_0_−g_i_)/g_0_* is the effect of drug *i* on the normalized growth rate *g/g*_0_ (Fig. 1; when effects are measured based on total yield rather than growth rate, Bliss additivity becomes multiplicativity; Supplementary Note 1). In contrast, according to Loewe additivity, the effect of drugs in combination is determined not by the sum of their normalized effects, but rather by the sum of their normalized dosages, such that their combined effect is the same across all combinations that have the same total normalized dosage. Namely, according to Loewe, lines of equal combination effect in drug-dosage space (isoboloes), are linear ^26^ (Fig. 1). For example, if two drugs are additive with respect to Loewe, their 50% inhibition isobole is a straight line satisfying *d*_1_/*d*_1_^50^ + *d*_2_/*d*_2_^50^ = 1, where *d_i_* is the dose of drug *i* and 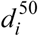 is the dose at which drug *i* alone causes 50% growth inhibition (IC50). Though conceptually different from one another, mechanistic support is available for both the Bliss and the Loewe models ^31^ and there is no agreement on which model should generally be used ^32–35^. Models that implement pairwise interaction data as well as higher order interactions can improve multi-drug predictions of either Bliss or Loewe ^21,36–39^. Yet, regardless of pairwise interactions, it remains unknown which of these two null models best predict the combined effect of multiple drugs.

**Figure 1:**
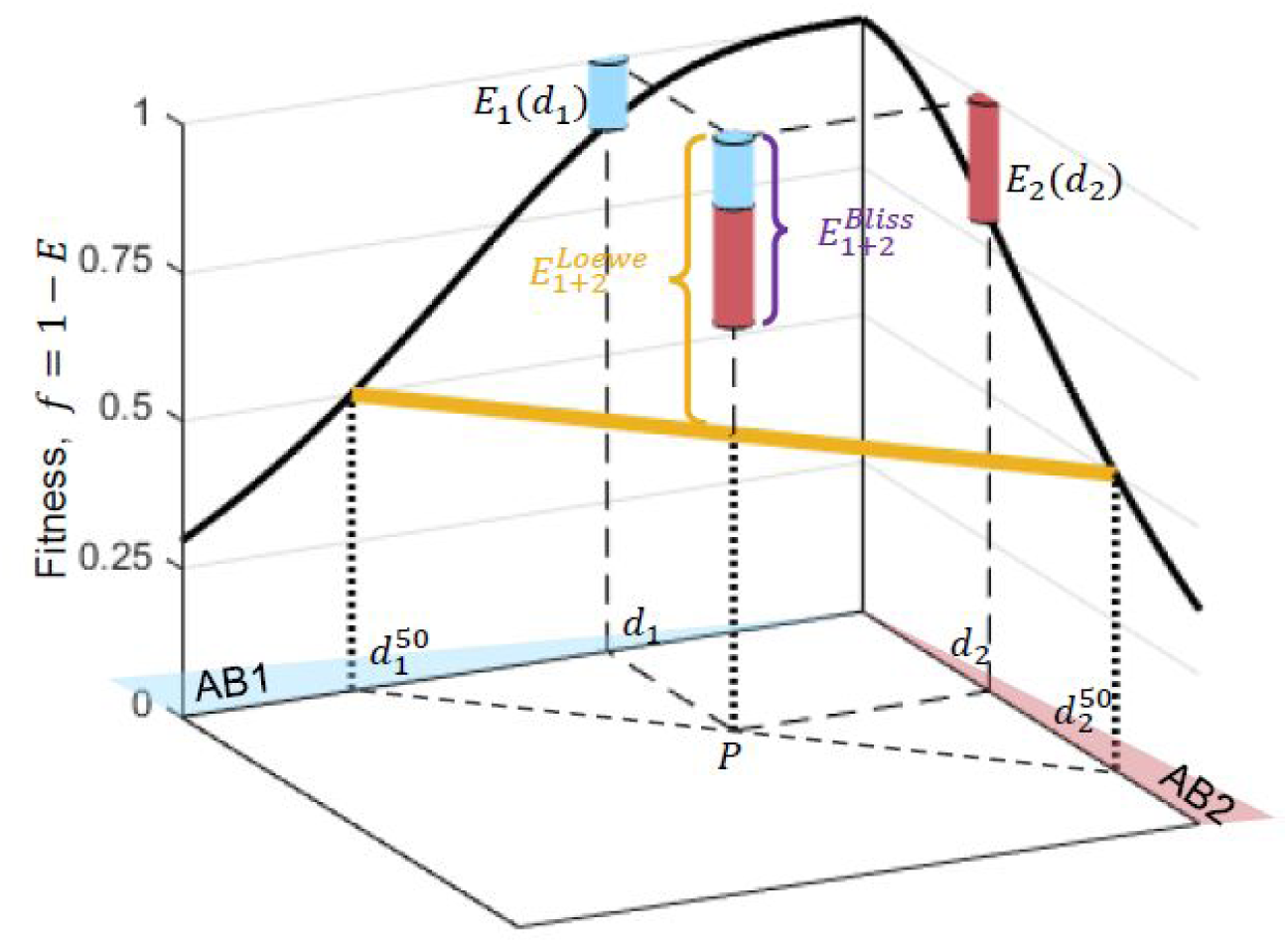
Schematic depiction of effect-additivity (Bliss) and dosage-additivity (Loewe). Given fitness as a function of dosage of each of the individual drugs (*f_i_* = 1 − *E_i_*, dose response curves, black solid lines), Bliss and Loewe models predict the fitness *f*_1+2_ = 1 − *E*_1+2_ at any given point *P* = [*d*_1,_ *d*_2_] in the drug concentration space. The Bliss prediction assumes additivity of normalized drug effects, 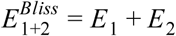, where *E*_1_ and *E*_2_ are the individual drug effects at their cognate concentrations (cyan and red piles, respectively). The Loewe model, on the other hand, assumes additivity of normalized drug dosage, such that the combined drug effect is fixed along linear lines of constant total normalized dosage (yellow, 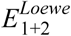 equals 50% in the example point *P*, and more generally is given by solving for 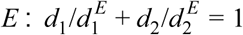, where 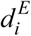 is the concentration of drug *i* that leads to inhibition level *E*).

Here, measuring bacterial response to antibiotic combinations, we contrast the Bliss and Loewe models for an increasing number of drugs, where we show that expectations of these models increasingly diverge. Quantitating bacterial response to combinations of up to 10 different drugs, we find that the Bliss model maintains accuracy with increased number of drugs, while the Loewe model loses its predictive power. Indeed, in contrast to Loewe, which predicts that the total drug dosage required for inhibition is constant, we find that total dosage increases monotonically with the number of drugs. Interestingly, our data show that this increase follows a square-root scaling, inspiring a simple model for orthogonality of dose additivity which follows a classical evolutionary optimization principle developed by R. A. Fisher ^12^.

To contrast the Bliss and Loewe models, we calculated how their predictions scale with increased number of drugs. As a natural measure of the combined potency of multiple drugs, we considered the total drug dosage needed to achieve a given level of inhibition. Defining “total dosage” (*D*) as the sum of the concentration of the individual drugs weighted by their corresponding IC50’s, 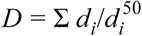, the “combination potency” (*D*^50^) is the total dosage *D* that yields 50% growth inhibition. As the number of drugs *N* increases, Loewe additivity predicts that the combination potency remains fixed, 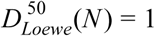. The prediction of Bliss, on the other hand, depends on the individual dose response curves of the different drugs. Assuming a Hill equation ^40^ for the single drug dose response 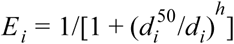, where *h* is the Hill coefficient, and equating the Bliss prediction of the combined effect 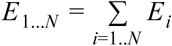 to 50%, yields the Bliss predicted scaling of combination potency with the number of drugs: 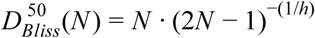. Thus, while Loewe predicts that the total dosage required for inhibition is constant, Bliss predicts that it increases with the number of drugs. The two models can therefore best be contrasted by measuring the combined action of increased number of drugs.

We considered 10 mechanistically different antibiotics and measured their effects on the growth rate of *Escherichia coli* populations, individually as well as in combinations of increased number of drugs. We chose antibiotics acting on a range of cellular functions, including cell wall synthesis, DNA replication, transcription and translation (Table 1). Measuring optical density (OD) versus time of bacterial growth on gradients of each of the individual drugs, we determined the dose response curve *g*(*d_i_*) for each of the drugs (Fig. 2a, Extended Data Figs 1–4, Extended Data Tables 1 and 2). These dose response curves are well fit by Hill functions, with Hill coefficients of the individual drugs ranging from 1 to 6.9 (Extended Data Figs 5 and 6). These fits also define the concentrations 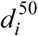 of 50% inhibition for each of the drugs in isolation.

**Table 1.** List of antibiotic used in the study

| # | Antibiotic | Abr. | Cellular Process | Target |
| --- | --- | --- | --- | --- |
| 1 | Rifampicin | RIF | RNA synthesis | RNA polymerase |
| 2 | Tetracycline | TET | Translation | 30S subunit of rRNA |
| 3 | Chloramphenicol | CHL | Translation | 23S subunit of rRNA |
| 4 | Erythromycin | ERY | Translation | 50s subunit of rRNA |
| 5 | Fusaric acid | FUS | Unknown | Dopamine beta-hydroxylase |
| 6 | Ciprofloxacin | CIP | DNA Synthesis | Gyrase |
| 7 | Cefoxitin | CEF | Cell Wall | PBP |
| 8 | Sulfamethoxazole | SMX | DNA Synthesis | DHPS |
| 9 | Trimetoprim | TMP | DNA Synthesis | DHFR |
| 10 | Phleomycin | PHL | DNA ds Breaks | DNA |

**Figure 2:**
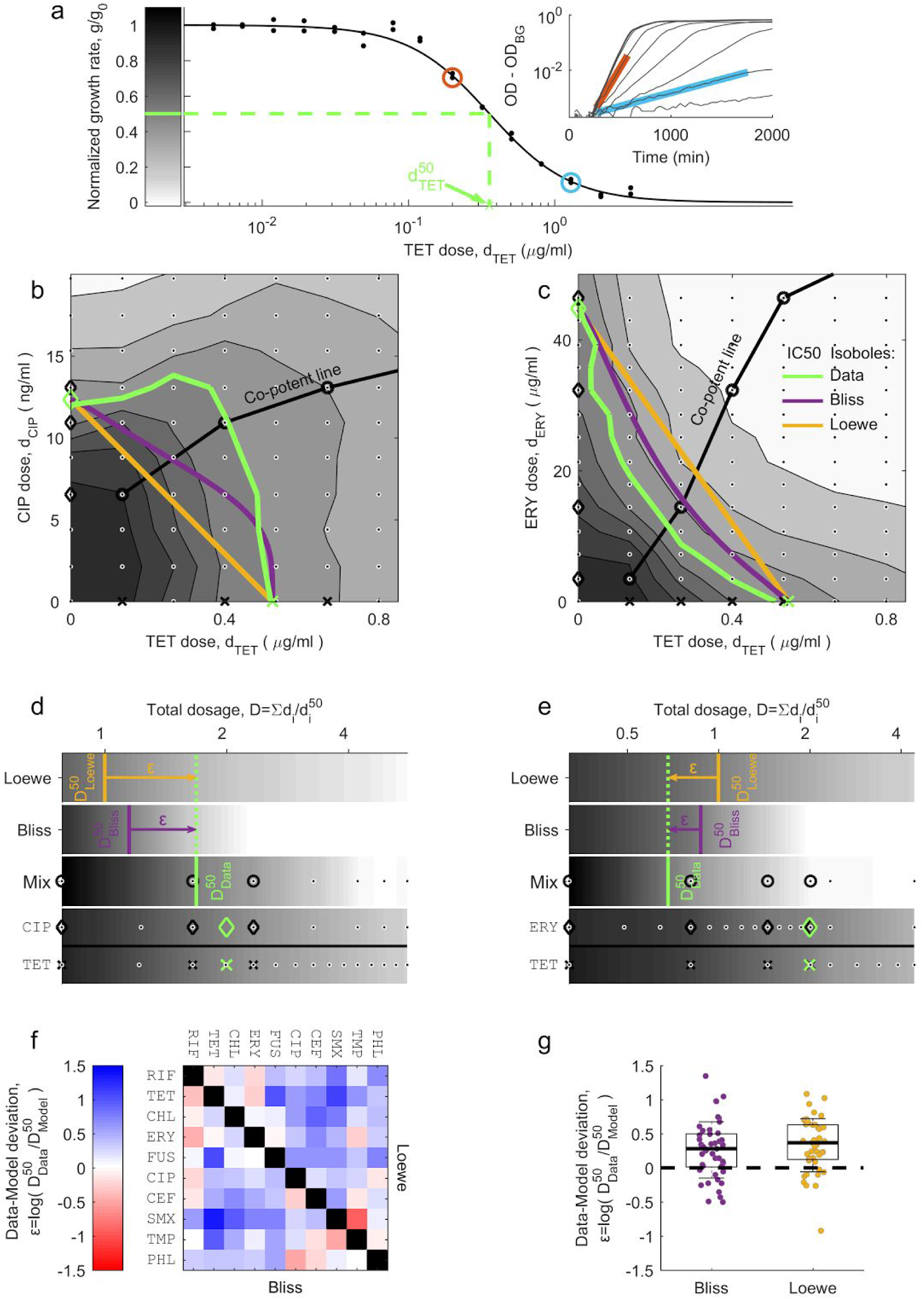
Pairwise measurements do not resolve the Bliss and Loewe models of additivity. **a**, Representing single drug dose response curve showing normalized growth rate *g/g*_0_ along a concentration gradient of TET (black dots, replicates), Hill equation fit (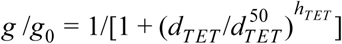, black line) and the IC50 (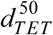, green dashed line). Inset: growth rates *g* were calculated by fitting OD_600_ measurements over time (black) to exponential function, *OD = OD_0_ · 2^g t^ + OD_BG_* (cyan and red). **b-c**, Response surface showing growth rates (grayscale indicated in panel A) over 2-D grid gradient (dots) of the antagonistic antibiotic pair TET-CIP (b) and the synergistic pair TET-ERY (c). The measured IC50 isoboles (green) are contrasted with Bliss (purple) and Loewe (orange) predictions. Indicated are the co-potent lines (circles), the corresponding co-potent single drugs (black x’s for TET, black diamonds for CIP and ERY), and the IC50’s (green symbols). **d-e**, Dose response along the co-potent line of the two drug mixtures (TET-CIP, d; TET-ERY, e) as a function of total dosage 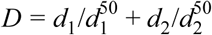. Measured normalized growth rates of the combined drugs (Mix) are contrasted with Bliss and Loewe predictions based on the single-drug data (shown below). Data-Model deviation, 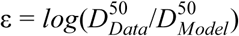, indicates the difference between measured 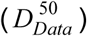 and predicted (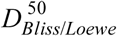) combination potencies. All symbols correspond to those in panels b-c. **f-g**, Data-Model deviations for each of the two models are presented as interaction matrix (f) and box plot (g). No significant difference between the models in their predictions of measured pairwise potencies (t-test, P=0.49).

Moving to drug pairs, we measured their combination potency and compared it to Bliss and Loewe predictions. We first measured the full response surface across 2-D dose gradients for two drug pairs: Tetracycline and Ciprofloxacin (TET-CIP) and Tetracycline and Erythromycin (TET-ERY), representing well-known examples of antibiotic antagonism and synergy (Figs 2b and 2c, response surface and IC50 isoboles) ^41,42^. Using the growth measurements of the individual drug gradients *E_i_*(*d_i_*), we derived the response surface and the IC50 isobole predictions of Loewe (straight line connecting the points 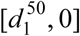 and 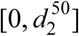) and Bliss (the set of all points [*d_1_,d_2_*] satisfying *E*_1_(*d*_1_)+ *E_2_*(*d_2_*) = 50%, Methods). As expected, the measured IC50 isobole lie above these predictions for the TET-CIP pair (indicating antagonism) and below for the TET-ERY pair (indicating synergy). While these two-dimensional gradients allow clear definition of synergy and antagonism, they require many growth measurements and become combinatorially prohibitive in a high-dimensional multi-drug space.

To effectively sample the concentrations space of multiple drugs, we performed growth measurements along a “co-potent” line ^38^, a curve in concentration space where the individual drugs have equal potencies in isolation (*E_1_*(*d_1_*) = *E_2_*(*d_2_*) = *… = E_N_* (*d_N_*), Figs 2b-e). This co-potent line sampling method vastly reduces the dimensionality of required measurements while guaranteeing that null models are evaluated in a region in drug concentration space where all drugs are active, rather than one in which the combined effect is dominated by a subset of drugs. Identifying the point *P =*(*d*_1_*, d*_2_,…, *d_N_*) on the co-potent line where growth is inhibited by 50% yields the combination potency, 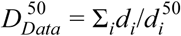. This measured combination potency was contrasted with the expected potencies 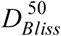 and 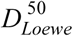, defined as the points along the co-potent line where the single-drug based calculations of Bliss and Loewe predict 50% inhibition (Supplementary Methods). The interaction between drugs was then defined as the deviation between the observed and predicted potencies of the combination 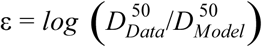, which captures the extent of antagonism (ε>0) and synergy (ε<0) (Figs 2d and 2e).

Measuring combination effects for all drug pairs, we find that their joint potencies are similarly well-predicted by both the Bliss and the Loewe models. For each of the 45 drug pairs, we measured their dose response along co-potent concentration gradient and determined their combination potency, 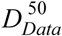 (supplementary methods, Fig. 2f, Extended Data Fig. 4). Comparing these combination potencies with predictions of the Bliss and Loewe null models, we find that both positive (ε > 0, antagonism) and negative (ε < 0, synergism) deviations are prevalent with respect to either model (Figs 2f and 2g). This prevalence of both antagonism and synergy among drug pairs overwhelms any small deviations between the two models (Fig. 2g; σ(ε*_Bliss_*)= 0.40; *σ*(*ε_Loewe_*) = 0.41; < ε*_Bliss_* >−< ε*_Loewe_* > =0.06, t-test: P=0.49). Further, clustering drugs based on these pairwise interactions, defined with respect to either Bliss or Loewe, leads to similar grouping by mechanism of action (Extended Data Fig. 7; possible small advantage to Bliss in resolving fine functionality differences) ^43^. The similarity of pairwise null predictions, the prevalence and magnitude of pairwise interactions with respect to both models, and their similar correlation with cellular function, prohibit distinction of the Loewe and Bliss null models based on drug pairs.

However, for increasing numbers of drugs, we find that their combined effect is well predicted by Bliss, while the Loewe prediction systematically diverges. Given that predictions of the two models should diverge with increased number of drugs, we measured the combined effect of multiple combinations with a varying number of antibiotics. We chose 35 combinations of three to ten antibiotics, including 8 randomly chosen sets from each combination size of *N* = 3, 5 and 7 drugs, all 10 sets of 9 drugs, and the whole 10-drug set (Figs 3a and 3b, Extended Data Fig. 8). Following the procedure used for the drug pairs, we measured the combined effect of each multi-drug set as a function of total dosage along co-potent lines and identified their combination potencies *D*^50^. Contrasting these measured potencies with the predicted potencies of Bliss and Loewe based on the single-drug measurements, we find that the Bliss model maintain good accuracy regardless of the number of drugs, while the accuracy of Loewe model declines as the number of drugs increases (Fig. 3c). The multiplicative form of the Bliss model (more suitable for yield measurements, Supplementary Note 1) is less predictive than the additive form, yet still much better compared to Loewe (Extended Data Fig. 9). We conclude that the Loewe model, predicting that the total dosage required for inhibition is fixed regardless of the number of drugs, can be rejected as a general predictor for the potency of multi-drug combinations.

**Figure 3:**
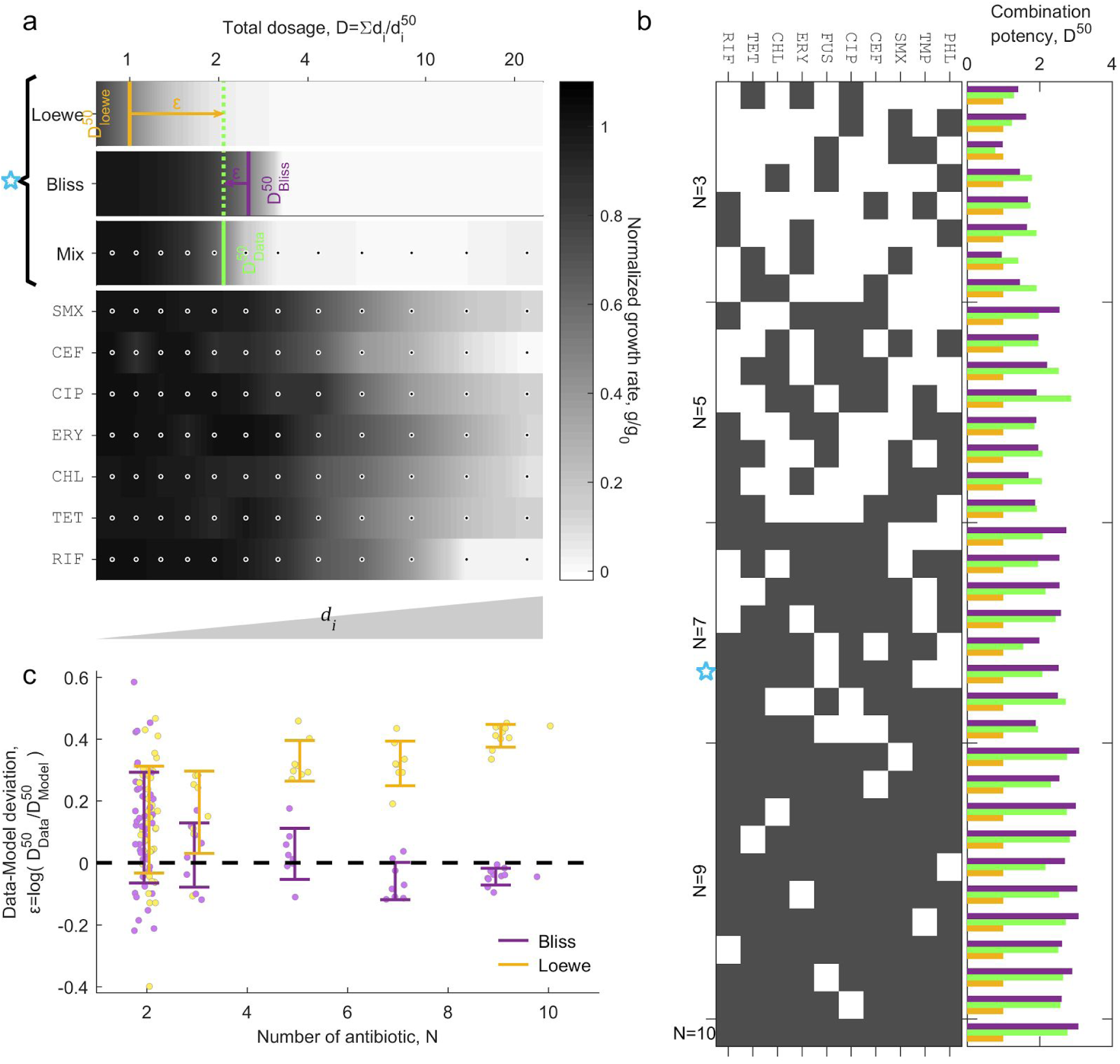
Loewe model of additivity loses its predictive power with increased number of combined drugs. **a**, An example of dose response along co-potent line of a mixture of 7 drugs. Measured normalized growth rate (gray scale) and combined potency 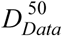 of the Mix are contrasted with Bliss and Loewe predictions calculated based the single drug measurements (below). **b**, Combination potency (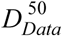, green) is contrasted with predictions of Bliss (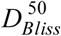, purple) and Loewe (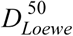, yellow) for 35 different combinations of 3 to 10 drugs (black squares). **c**, Deviation of each of the models from the data 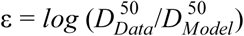 is plotted as a function of number of drugs, showing that the Loewe predictions deviate from the data with increased number of drugs, while Bliss predictions remain accurate.

Next, we tested how the potency of drug combinations, namely the total dosage required for inhibition, varies with the number of drugs. To account for any slight experimental deviations from the ideal co-potent line, we use a natural entropy-like definition of an effective number of drugs *N_eff_* which is based on the uniformity of the individual drug effects *(N_eff_* equals *N* if all drugs have the same effect; is slightly smaller than *N* when these effects are uneven; and converges to 1 at the extreme case when a single drug dominates and all the other have no effect; see definition of *N_eff_* in Fig. 4 caption). Contrary to the Loewe prediction, we find that the total dosage required for inhibition increases with the effective number of drugs (Fig. 4a). Moreover, this inhibitory total dosage seems to obey a simple scaling law: it increases as the square root of the effective number of drugs (*D*^50^ = (*N_eff_*)*^α^*, least square fit yields: *α* = 0.48 ±0.03).

**Figure 4:**
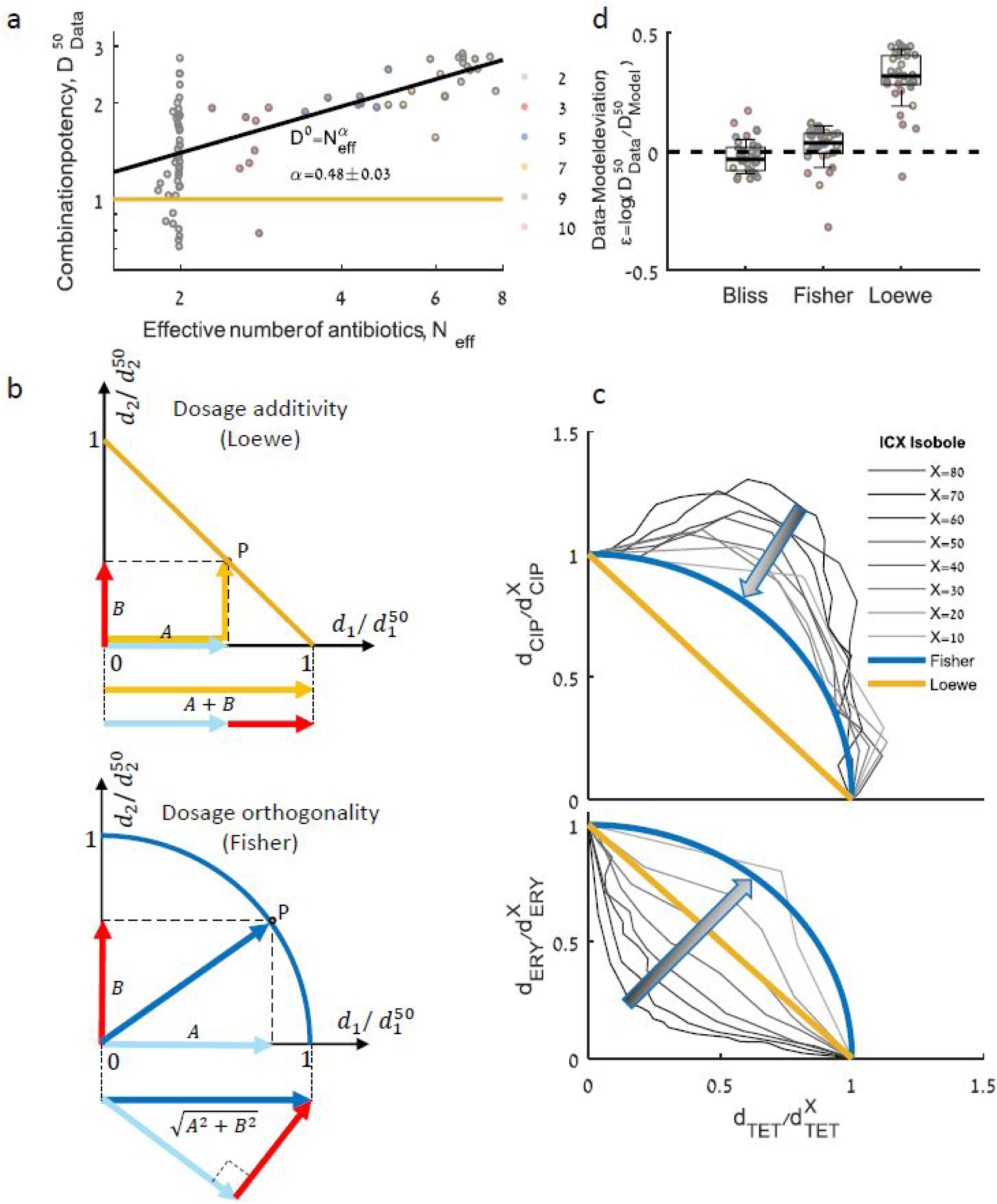
A square-root scaling law of inhibitory total dosage with effective number of drugs is explained by a simple dosage-orthogonality model. **a**, Combination potency, 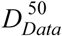 of all 80 different drug combinations is plotted as a function of effective number of drugs 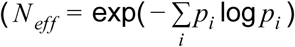, where 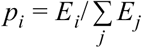, and *E_i_(d_i_)* are the single drug individual effects at their cognate concentrations; colors represent the actual number of drugs, *N*). In contrast to Loewe, which assumes that the total dosage required for inhibition is fixed (yellow line), the total dosage increases as square root of the effective number of drugs (black line, fit of *D*^50^ = (*N_eff_*)*^α^* yields α = 0.49 ±0.03, 0.95 confidence interval). **b**, The square-root scaling is explained by a Fisher inspired dose-orthogonality model, which assumes that for small perturbations the dosages of independent drugs should be added as orthogonal vectors rather than linearly as in Loewe. Hence isoboles of *X%* inhibition are spherical surfaces defined by 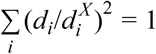 (Fisher, bottom, circles in two-drug space, blue line) instead of linear surfaces (Loewe, top, straight lines in two-drug space, yellow line). **c**, Indeed, even for strongly interacting drug pairs CIP-TET (top) and ERY-TET (bottom), isoboles (grey lines) become more circular (Fisher prediction, blue) rather than linear (Loewe prediction, yellow) as the inhibitory effect X approaches zero. **d**, Data-model deviation 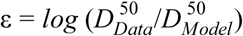 of all the combinations of more than two drugs for each of the three models. The data strongly reject Loewe and is instead consistent with both Fisher and Bliss (t-test: Loewe, P<10^−16^; Bliss, P=0.26; Fisher, P=0.1).

The square root scaling of the inhibitory dosage with number of drugs can be explained by an approximation to Bliss, inspired by the classical optimization principle of Fisher’s geometric model of adaptation ^12^. Fisher’s model describes the fitness *f* in a space of *N* independent orthogonal phenotypes and assumes that it declines as a function of the euclidean distance from an optimal point (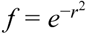, where 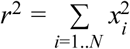 and *x_i_* are phenotypic distances from the optimal point). For a given fitness value, the phenotypic distances *x_i_* therefore decline as 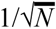 and their sum, 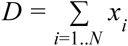, increases as 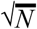. The analogy of drug dosages with Fisher’s phenotypes explains the square root scaling of total inhibitory dosage with number of drugs and underscores that drug concentrations should be summed not linearly by simple addition as in Loewe, but rather as the geometric sum of orthogonal vectors (Fig. 4b; Of course, orthogonality is a null idealization from which drug combinations can deviate due to synergy or antagonism, or when similar drugs act along the same axis). This Fisher’s inspired “dose-orthogonality” model can also be derived as an approximation of Bliss additivity at the limit of small dosages (Supplementary Note 2). Indeed, we find that even strongly interacting drug pairs assume more circular isoboles for small fitness effects (Figs 2b, 2c, 4c). Similarly to Bliss and in contrast to Loewe, combination potency predictions of the dose-othethognality model (derived by intersecting the co-potent lines with spherical IC50 isoboloes 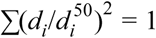, Methods), are consistent with the drug combination measurements (Fig. 4d). Yet, unlike Bliss these predictions do not require fine measurements of the minute individual drug effects *E_i_(d_i_*), but rather depend on the more robust measurements of their individual IC50 dosages, 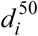. Using Loewe’s dose language, the dose-orthogonality model provides an intuitive and robust approximation of Bliss (Extended Data Table 3), which predict the potency of drug combinations similarly well and explain the square-root scaling law of potency with number of drugs.

Our measurements reject the Loewe model of dosage-additivity for predicting combination of multiple diverse drugs, favoring the Bliss effect-additivity and motivating a simple model of dosage-orthogonality. In contrast to Loewe additivity, which predicts that the total dosage required for inhibition is fixed, we find that the total inhibitory dosage increases with the number of drugs, following a square root scaling law. This general reduction in potency with increased number of drugs implies that bacterial inhibition by multi-drug combinations may be require higher total drug dosages than classically anticipated. The square root scaling supports a model for drug additivity where dosages of independent drugs add up orthogonally rather than linearly. This dosage-orthogonality model provides an approximation to Bliss, yet it uses dosage arithmetics which allows a more robust implementation and simple intuition. It will be interesting to explore the generality of these results and the limit on the number of orthogonal pharmacological axes as more antibiotics and stresses are added, as well as beyond the minimal inhibitory concentration and in more complex systems such as in cancer therapy. Throughout such clinical systems and natural ecologies, our findings provide a uniform framework for understanding the null arithmetics of many-drug combinations.

